# Understanding Others Requires Right Temporoparietal and Inferior Frontal Regions

**DOI:** 10.1101/2021.03.31.437941

**Authors:** Tatiana T. Schnur, Junhua Ding, Margaret Blake

## Abstract

The human ability to infer other people’s knowledge and beliefs, known as ‘theory of mind’, is an essential component of social interactions. Theory of mind tasks activate frontal and temporoparietal regions of cortex in fMRI studies. However, it is unknown whether these regions are critical. We examined this question using multivariate voxel-based lesion symptom mapping in 22 patients with acute right hemisphere brain damage. Studies of acute patients eliminate questions of recovery and reorganization that plague long-term studies of lesioned patients. Damage to temporoparietal and inferior frontal regions impaired thinking about others’ perspectives. This impairment held even after adjustment for overall extent of brain damage and language comprehension, memory, comprehension, and attention abilities. These results provide evidence that right temporoparietal and inferior frontal regions are necessary for the human ability to reason about the knowledge and beliefs of others.

## 1.0 Introduction

An essential component of social communication is the human ability to understand that others have beliefs and knowledge that differ from one’s own, known as theory of mind (ToM). Functional imaging evidence suggests a network of brain regions support ToM. Yet, which regions are necessary for ToM is unknown as lesion data come from participants with large chronic lesions which obscures anatomical specificity. Critically, reorganization during recovery may have shifted function to other regions^1–4^. Here we address these critical gaps in a rarely studied clinical population of adults with focal acute right hemisphere stroke to determine which regions are necessary for ToM.

ToM is critical for social communication. Discourse comprehension requires understanding what a speaker means, especially when in opposition to what was actually said, such as occurs in sarcasm, humor, idioms and metaphors^5–13^. Human communication is rife with nonliteral or indirect language and meanings that must be deduced by considering the speaker’s perspective. Inappropriately responding during conversation to misunderstandings of what a speaker knows and intends creates increasingly poor social interactions and relationships^13–15^.

Distressingly, communication deficits have profoundly damaging effects on quality of life by decreasing return to work, social participation, and interpersonal relationships^5,16–23^ which result in increased morbidity and mortality^24,25^. Thus, uncovering the behavioral and neural mechanisms underlying right hemisphere (RH) deficits will help understand their impact on quality of life with the goal of developing better methods for assessment and intervention thus improving access to health care and long-term outcomes for this underserved patient population^5,26,27^.

First-order ToM, knowing what a protagonist knows, is typically unaffected by RH stroke^28,29^. However, second-order ToM, inferring what a protagonist knows about *another* character’s knowledge or beliefs is impaired after RH stroke^5,19,30,31^. Individuals with RH stroke exhibit dissociations in implicitly thinking about other peoples’ perspectives, referred hereafter as *Other* ToM^32–36 37,38^ and explicitly managing other people’s perspective separate from their own, referred to as *Self* ToM ^32,33,36,37,39^. *Other* and *Self* ToM vary in the degree of ToM-related inhibition^40^, but critically, are independent of general inhibitory/executive functions^39^.

ToM is subserved by a large ‘mentalizing’ network involving right anterior brain regions (medial to dorsal to ventral-lateral prefrontal cortex, the precentral gyrus, posterior cingulate cortex, and the insula) and bilateral posterior brain regions (the temporal-parietal junction, TPJ)^19,41–49^. Neuropsychological studies which have examined the necessity of these brain regions for ToM function have identified the right inferior frontal gyrus (rIFG), but not rTPJ regions as critical for explicit *Self* ToM^39,50,51^ while others found that damage to both rTPJ and rIFG led to poor *Self* ToM performance^33,36,37^. Implicit *Other* ToM processing deficits have been related to damage to left and/or right TPJ regions but not rIFG regions^32,34–36^. The discrepancies across patient studies likely derives from multiple cross-experiment differences including the use of different tasks to assess *Other* and *Self* ToM and differing patient selection criteria based on lesion location^32,34,35,39,52^ and/or deficit severity^36,37,39^. Results are further confounded by a lack of control for overall lesion volume and task demands^37,51^ and most studies examine these questions in individuals during the chronic stage of stroke, after reorganization of function has likely occurred. Thus, it is unknown whether damage to this network of regions is uniquely related to ToM deficits, related to overall stroke severity, cognitive dysfunction related to baseline task demands, or reorganization of function.

### 1.1 Current Study

The purpose of this study was to identify the brain regions necessary for *Self* perspective and *Other* perspective ToM before reorganization of function in adults with acute RH stroke. We used a visual non-verbal false belief (FB) task to assess two subcomponents of ToM, the ability to implicitly and spontaneously infer another person’s perspective (*Other* ToM), and the ability to explicitly manage conflict between self-perspective and another person’s perspective (*Self* ToM)^33,34,40^. We chose this ToM assessment for several reasons. First, the task is non-verbal, reducing the confound that poor performance on ToM is due to a language impairment^53,54^. Second, all trials involve the same scenario (in contrast with other story-scenario ToM tasks^30,31,55^). Attention, memory, executive functioning, and language input processing demands are consistent across ToM subcomponent trials, thus controlling confounds that could otherwise contribute to performance differences between ToM subcomponents^56,57^. Third, the task is a standardized diagnostic procedure. Biervoye et al.^40^ demonstrated no effects of gender or socio-economic status on ToM. Fourth, the task includes interleaved control trials to assess the contribution of working memory, attention, and comprehension, independent of ToM ability while controlling for unilateral neglect. Thus, we can understand the degree to which ToM difficulties result from a general cognitive deficit. Fifth, the task has been implemented across different clinical populations (stroke^32,34,40^; acquired brain injury^40^; alcoholism^58^; schizophrenia^59^; and elderly subjects^40,60^) demonstrating task feasibility across populations. Lastly, *Other* and *Self* ToM as measured in this task have been shown to dissociate from domain general inhibition processes^39^.

## 2.0 Methods

### 2.1 Subjects

We consecutively recruited 37 subjects who presented with neurological signs of an acute right hemisphere stroke from three comprehensive stroke centers in the south-central United States. Thirteen subjects were subsequently excluded either because of no neuroradiological signs of acute stroke (n=1), previous stroke or traumatic brain injury (n=3), hemianopia (n=2), history of drug abuse (n=2), hospital discharge precluding test administration (n=2), or an inability to complete the task battery (n=3). The 24 subjects included in the analyses (8 Female) were native English speakers and had no history of other significant neurological disease. Average age and education were 58.6 years (SD 13.6; range 25 – 82) and 12.3 years (SD 2.6; range 5 – 19; 1 subject unknown), respectively. The Baylor College of Medicine Institutional Review Board approved the informed consent. Subjects received financial compensation for participation.

### 2.2 Design

Subjects completed behavioral testing at bedside within an average of 2.8 days after stroke onset (SD 1.6; range 0 – 6 days). We presented stimuli via laptop (when appropriate) and recorded responses by digital recording and by hand where possible.

### 2.3 Baseline cognitive and language tasks

#### 2.3.1 Unilateral neglect

To assess the presence of unilateral neglect we administered the apples cancellation task^61^from the Birmingham Cognitive Screening^62^ and an adaptation of the Ogden scene copy task^63^. Both tasks measure egocentric (viewer-centered) and allocentric (stimulus-centered) neglect. The dependent variable was the average percent correct of right vs. left sided stimuli for both egocentric and allocentric neglect across both tasks.

#### 2.3.2 Working memory

To assess working memory ability, we administered the digit span subtest from the Repeatable Battery for the Assessment of Neuropsychological Status^64^. Subjects heard sequences of two to nine numbers at approximately one word per second (two sequences per list length), after which they were asked to repeat the sequence back in the correct order. There were two sequences per list length. The second sequence was administered only when participants failed the first one. We discontinued the task after failure of both sequences in any list length. The dependent measure was the estimated span length for 75% correct, using linear interpolation between the two list lengths that span 75%.

#### 2.3.3 Language comprehension and production

To rule out frank aphasia, single word comprehension was assessed through shortened versions of word-picture matching (17 pictures paired with matching, phonologically related, and semantically related words^65^) and synonymy triples (n=12)^66^. Naming was assessed with a short form of the Philadelphia Naming Battery (n=30)^67^. See Table 1 for average performance and range across baseline cognitive and language tasks.

**Table 1.** Summary of baseline task performance including measures of neglect, working memory, word comprehension and production.

| Task | M (SD) | Range |
| --- | --- | --- |
| Neglect – egocentric | 12% (.25) | -24 – 83% |
| Neglect – allocentric | 2% (.10) | -17 – 27% |
| Working memory | 5.6 (1.1) | 3.3 – 8.3 |
| Word-picture matching | 92.4% (5.2) | 84.3-100 |
| Synonymy triples | 87.3% (10.9) | 58.3-100 |
| Picture naming | 96.3% (5.0) | 83.3-100 |

N.B. One subject was unable to complete the word-picture matching, synonymy triples, and picture naming tasks.

### 2.4 Theory of mind assessment

We used a visual non-verbal false belief task to assess two subcomponents of second-order ToM^40^. The first component, *Other* (perspective) ToM assesses the ability to spontaneously infer another person’s perspective without explicit instruction to do so. The second component, *Self* (perspective) ToM assesses the ability to explicitly infer another person’s perspective which requires managing conflict between self-perspective and another person’s perspective. Similar to classic ToM tasks (for a review see Martín-Rodríguez & León-Carrión, 2010)^68^, the Biervoye et al.^40^’s assessment uses scenarios in which the viewer has more information than a character in the scene and must decide what the character knows. Subjects see a series of short video trials with no sound. For example, a man places a green object in one of two boxes in sight of a woman. She then leaves the room and when she returns, she indicates where the object is. The experimenter explained that the woman’s role is to help the subject identify the object location. Conditions differ in the following respects. In all False Belief Trials (whether *Other* or *Self* ToM) the man switches the location of the two boxes. However, in the *Other* ToM False Belief condition, the subject does not see where the object was initially placed. At the end of *Other* ToM trials, the subject answers the question “show me where the green object is”. For correct False Belief trial responses, the subject must understand that the woman has a FB about the object location because the boxes were switched during her absence. As a result, the woman put the marker on the wrong box and the object must be in the other box. Because the subject did not see where the object was placed originally, subjects must infer the woman had a false belief before knowing the location of the object. As a result, subjects do not have especially strong self-perspective competing knowledge about the location. In contrast, in the *Self* ToM condition the subject sees where the object is hidden initially. Subjects are asked to point to the box that the woman will open first to find the green object. Thus, when asked where the woman will look for the object, they must manage conflict with their own knowledge to respond using only the woman’s knowledge.

Interleaved control trials assessed the contribution of working memory, attention, and comprehension, independent of ToM ability while controlling for unilateral neglect. Memory control trials assess working memory where the woman indicates the object location before the object location switches. This requires the subject to hold the object location in working memory and update it when it switches at the end of the trial. In filler trials, part of the control condition, subjects see that the object is switched in the presence of the woman (by which the woman can then correctly indicate the location of the object). Poor performance on filler trials demonstrates a deficit in some aspect of input processing beyond any potential deficit in ToM reasoning. Lastly, in the True Belief control trials any manipulation of the object (e.g., lifting a box rather than switching boxes) never changes the object’s location. If a subject makes more errors on the True Belief trials (when the location never changes) compared to the FB trials this suggests that the subject used a strategy in the FB trials such as picking the box which the woman did not point at. See Biervoye et al.^40^ and Samson, Apperly, Chiavarino, and Humphreys^69^ for further details and rationale concerning trial types.

#### 2.4.1 Procedure

We followed task administration as outlined in Biervoye et al.^40^. To summarize, we presented 48 trials divided into *Other* ToM and *Self* ToM conditions in pseudo-random order. We presented the *Other* ToM condition always first to keep the demands of the *Self* ToM high (cf. Biervoye et al.^40^). We provided task instructions and separate practice sessions before each condition. Each condition included False Belief trials (*Other* ToM, n=8; *Self* ToM n=8) and Control trials (*Other* ToM n=12, *Self* ToM n=16). We normalized subjects’ performance across trial type in comparison to age-matched scores provided by Biervoye et al.^40^ (p. 11, Table 5)^1^.

### 2.5 Imaging acquisition and lesion tracing

Nineteen participants received MRI and five participants received CT imaging, all acquired in the axial direction. Neuroimaging was collected within an average of two days of behavioral testing (range 0-5 days). For those with MRI imaging, their DWI images (1 * 1 * 5 mm) were first registered to higher resolution structural images (T1/T2 FLAIR; 0.5 * 0.5 * 5 mm) using AFNI^70^ (https://afni.nimh.nih.gov). Then participants’ lesions were demarcated on DWI images using ITK-SNAP^71^ (https://www.itksnap.org). Finally, structural images and lesion masks were normalized into MNI space using ANTs^72^ (https://stnava.github.io/ANTs/). For those with CT images (0.5 * 0.5 * 5 mm), lesions were directly demarcated on the Colin-27 template. For two subjects, lesions were not clearly identifiable on the CT scans and thus they were excluded from lesion-symptom mapping analyses.

### 2.6 Multivariate lesion symptom mapping

Support vector regression lesion symptom mapping (SVR-LSM) was performed using LESYMAP^73^. We used epsilon-regression SVR with its default parameters (radial basis function kernel, gamma =5, cost = 30, epsilon = 1). We performed two SVR-LSMs to measure the relationship between brain damage location and either the residuals of the *Other* ToM and *Self* ToM normalized *z*-scores, while controlling for performance in each ToM’s respective control trials as well controlling for the contribution of total lesion volume (number of lesion mask voxels). We included only voxels with damage in at least two participants (>5%). Voxels were combined as patches if they had the same lesion pattern across all participants. The patches’ statistical inference was estimated by using 10000-times permutation on the dependent variable^74^. The significant regions (*p* < 0.05) were mapped onto published gray matter^75^ to report their locations.

## 3.0 Results

### 3.1 Behavior

To provide a clinical perspective as to the incidence of ToM deficits in the acute RH stroke population, we followed Biervoye et al.’s^40^ categorization procedure to assess subjects’ abilities to infer what a protagonist knows about another character’s knowledge or beliefs (spontaneous other-perspective processing, *Other* ToM) and abilities to represent other people’s perspective separate from their own (management of self-perspective, *Self* ToM). To assess whether *Other* and *Self* ToM processing were impaired because of the influence of input processing deficits, we measured the integrity of input processing (attention, memory, and language comprehension) via *Other* and *Self* associated Control trials. See Supplemental Table 1 for subject ToM behavior data and ToM classifications.

#### 3.1.2 *Other* ToM performance

As seen in Figure 1A, subjects demonstrated variability in the *Other* ToM and associated control condition (False Belief mean = −1.8, median = −1.7, range −4.7– 1; Control trial mean = − 2.9, median = −1.6, range −18.4 – .4). We classified subjects as having deficits in the *Other* ToM associated conditions (False Belief, Control) if subjects scored below 2 SDs from their age-matched control mean. Notably, 9/24 subjects performed below the control performance cut-off for the *Other* ToM False Belief trials (in Figure 1A see individuals in comparison to the red dotted line in the False Belief condition). We then compared performance between the False Belief and Control conditions to assess whether the FB deficit reflected a deficit beyond that observed on the control trials (a difference of > 2 SD). Of the nine subjects with an *Other* ToM False Belief deficit, five evidenced a significant dissociation between False Belief and Control Trials (and had spared input processing) where the other four subjects’ False Belief performance could not be distinguished from input processing difficulties.

**Figure 1.**
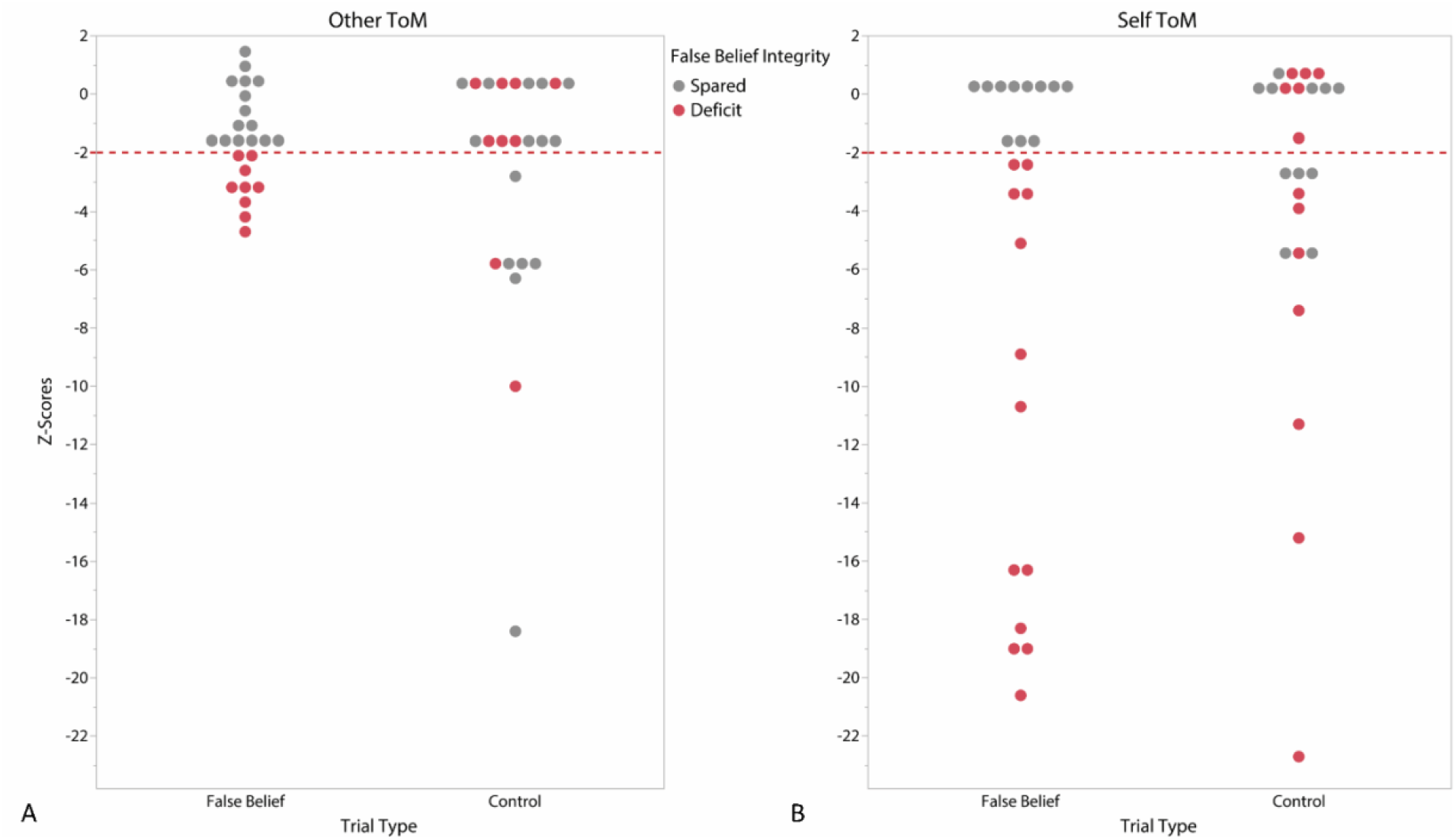
*Other* ToM (Figure 1A), *Self* ToM (Figure 1B) and Associated Control Trial Performance. The red-dashed lines indicate the −2 SDs cut-off thresholds across conditions below which performance was impaired in comparison to age-matched non-brain damaged subjects. Circles reflect individual z-scores. Red circles indicate individuals who performed below the cut-off threshold on the False Belief trials and grey circles indicate individuals who performed above the cut-off threshold.

In sum, of the 24 subjects tested during the acute phase of a first-onset right hemisphere stroke, 21% of subjects demonstrated an *Other* ToM processing deficit independent of any input processing difficulties and 62% of subjects demonstrated spared *Other* ToM processing. Assessment of *Other* ToM processing for the remaining 17% of subjects was obscured by input processing difficulties.

#### 3.1.3 *Self* ToM trial performance

We followed the same procedure as *Other* ToM to assess the clinical incidence of *Self* ToM deficits in comparison to control trials (False Belief, Control). As seen in Figure 1B, subjects demonstrated variability across *Self* ToM associated conditions (False Belief mean = − 6.2, median = −2.4, range −20.6 – .4; Control mean = −3.6, median = −2.1, range −22.7 – .5). We classified subjects as having deficits in the *Self* ToM associated conditions (False Belief, Control) if subjects scored below 2 SDs from their age-matched control mean. Notably, 13/24 subjects performed below the control performance cut-off for the *Self* ToM False Belief trials (in Figure 1B False Belief condition see individuals in comparison to the red dotted line). Comparing performance between the False Belief and Control conditions for the 13 subjects with a *Self* ToM False Belief deficit, 11 of 13 evidenced a significant dissociation between False Belief and Control Trials where the remaining two subjects’ False Belief performance could not be distinguished from input processing difficulties.

In sum, of the 24 subjects tested during the acute phase of a first-onset right hemisphere stroke, 46% of subjects demonstrated a *Self* ToM processing deficit independent of any input processing difficulties and 46% of subjects demonstrated spared *Self* ToM processing. Assessment of *Self* ToM processing in the remaining 8% of subjects was confounded by input processing difficulties.

#### 3.1.4 Direct comparison between *Other* and *Self* ToM performance

We quantified the number of subjects who showed dissociations between *Other* and *Self* ToM given a dissociation between *Other* and/or *Self* ToM and associated control trial performance. Three subjects overlapped between the five subjects who showed an *Other* ToM deficit (compared to control performance) and the 11 subjects who showed a *Self* ToM deficit (compared to control performance). Thus, we evaluated 13 different participants for a dissociation between *Other* and *Self* ToM. To do so, we compared the absolute difference in performance between each ToM task (*Other* – *Self* ToM) z-scored in comparison to age-matched controls. We considered differences of > 2 SDs reflective of a significant dissociation (either a classical dissociation of one ability impaired, the other intact or a strong dissociation where both ToM abilities were impaired) between *Other* and *Self* ToM processing (see Supplemental Table 1 for individual classifications).

Of the 13 subjects evaluated for a dissociation between *Other* and *Self* ToM, six subjects were impaired on *Self* ToM but with spared *Other* ToM and one subject was impaired on *Other* ToM but with spared *Self* ToM. Three subjects were impaired on both *Other* and *Self* ToM where two of the three demonstrated a strong dissociation (*Self* ToM more impaired than *Other* ToM). Three other subjects showed significant dissociations between FB performance (one subject was more impaired on *Other* ToM than *Self*, the other two subjects more impaired on *Self* vs. *Other* ToM). In these latter three cases, the dissociation may not be as interpretively clear-cut as subjects demonstrated input processing difficulties which obscured performance in the opposing FB condition.

#### 3.1.5 Summary

Overall, 54% of the 24 subjects demonstrated some form of ToM impairment independent of input processing difficulties during the acute stage of stroke. Of these, 8% were impaired only on *Other* ToM, 33% were impaired only on *Self* ToM, and 13% were impaired on both ToM types. Remaining subjects showed no significant FB deficits after considering the effect of input processing difficulties, where 33% showed intact processing across all conditions and for 13% FB trial performance was affected by input processing difficulties.

### 3.2 Lesion Symptom Mapping

#### 3.2.1 Lesion distribution

Figure 2 top panel illustrates the distribution of overlapping lesioned voxels damaged in at least two participants across 22 participants with acute RH damage (lesion size: M = 28030 mm^3^, S.D. = 46369 mm^3^, range = 279-194487 mm^3^). Overlapping lesion coverage was extensive and included the middle and inferior frontal gyri, orbital gyrus, pre- and post-central gyri, superior, middle and superior temporal gyri, superior and inferior parietal gyri, insular cortex, basal ganglia and thalamus.

**Figure 2.**
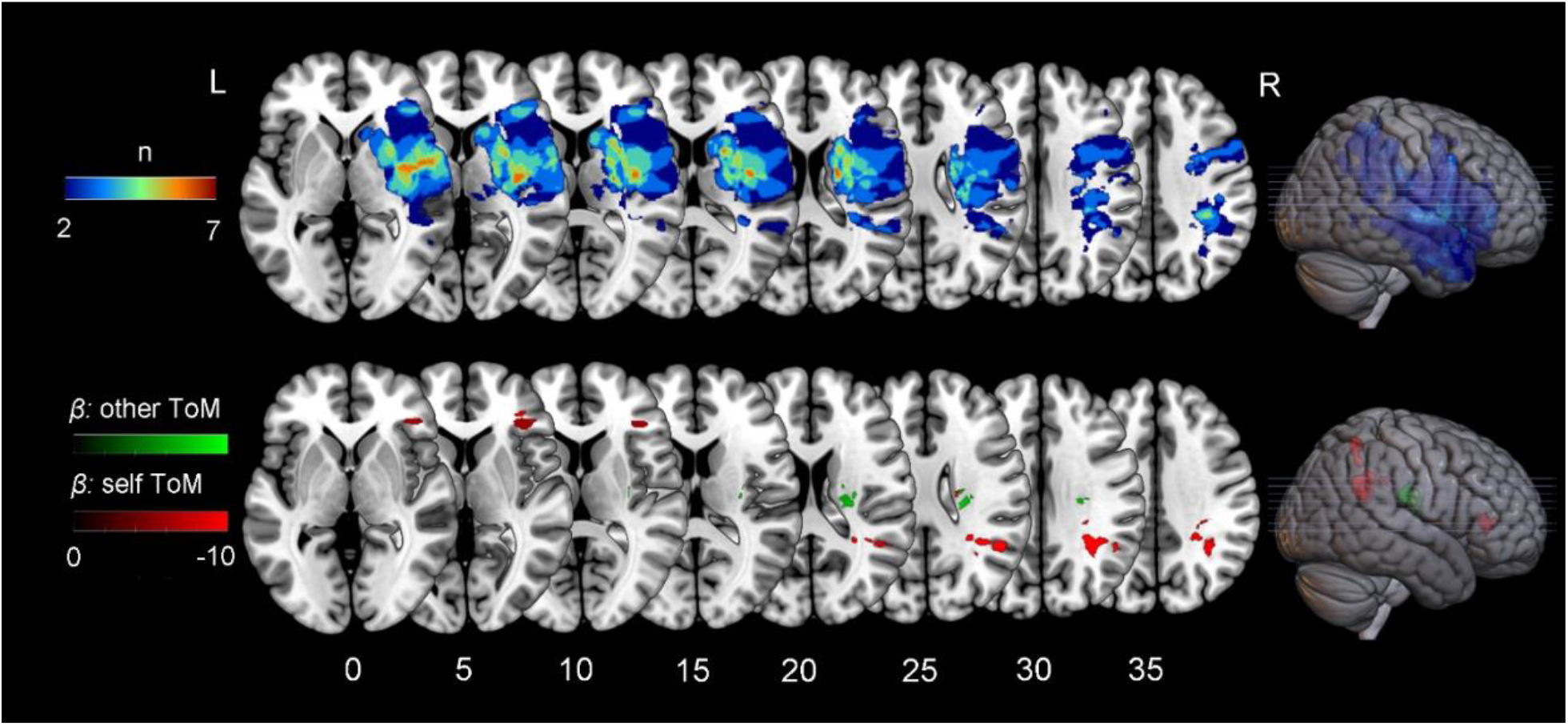
Lesion Overlap Distribution and SVR-LSM Results. Top panel illustrates overlapping (in at least two participants) acute RH damage across 22 participants included in the LSM analyses. Bottom panel illustrates location and beta values of lesioned voxels significantly related to impairments in *Other* ToM (green) or *Self* ToM (red), while controlling for the opposing ToM ability and total lesion volume. Axial slices are displayed from z coordinates 0-35 mm adjacent to RH surface projections.

#### 3.2.2 SVR-LSM of Theory of Mind impairments

Impairments in *Self* ToM were significantly associated with damage to the opercular region of the IFG (BA 44), the rostroventral inferior parietal lobe (BA 39) part of the angular gyrus within the dorsal posterior TPJ^76–78^, and a smaller region within the superior parietal lobule (BA 7). Results were independent of overall lesion volume and performance in the associated control condition requiring language comprehension, working memory, and attention. Other significant regions were sparse and involved fewer than 50 voxels. Impairments in *Other* ToM, independent of overall lesion volume and the control condition were associated with undifferentiated white matter damage. To note, we did not include demographic variables as regressors in the LSM analyses (i.e. age, number days tested after stroke onset and level of education) as none significantly correlated with either *Other* or *Self* ToM performance (*p*’s > 0.13). See Table 2 for a list of regions and number of voxels significant in the SVR-LSM analyses.

**Table 2.** Regions significantly associated with either *Self* (perspective) and/or *Other* (perspective) ToM.

| Lobule | Gyrus | Anatomical and Cyto-architectonic Descriptions | *Self* ToM (mm<sup>3</sup>) | *Other* ToM (mm<sup>3</sup>) |
| --- | --- | --- | --- | --- |
| Frontal Lobe | Inferior Frontal Gyrus | opercular BA 44 (opercular inferior frontal gyrus) | 258 | 0 |
| Parietal Lobe | Inferior Parietal Lobule | rostroventral BA 39 | 431 | 0 |
|  | Superior Parietal Lobule | Rostral BA 7 | 107 | 0 |

## 4.0 Discussion

Approximately half of the consecutively enrolled acute RH stroke cohort demonstrated impairments in explicit reasoning about others. Our LSM results demonstrate that damage to the rTPJ and rIFG impairs explicit reasoning about other’s beliefs when they differ from one’s own. The relationship between damage and explicit reasoning about others was not due to overall extent of brain damage, reorganization of function, or impairments in other cognitive abilities including language comprehension, memory, or attention. In contrast, we found a lower incidence of impairments in implicit reasoning about others and no relationship with damage to any right hemisphere gray matter region. Together these results demonstrate that the rTPJ and rIFG play a critical role in explicit mentalizing about others.

In the current study, over 50% of acute RH stroke participants had a deficit in at least one form of ToM that was not attributable to deficits in attention, working memory, or task complexity. The rate of incidence is less than the 68% reported by Balaban and colleagues^30^. However, their participants were recruited from rehabilitation centers and thus were more likely to have cognitive-communication disorders^79,80^ than a group selected only on lateralization of lesion. Additionally, Balaban and colleagues assessed ToM with a battery of eight different tasks and no participant was impaired on all tasks suggesting that with a smaller battery the rate of identification could be substantially lower. The incidence of ToM deficits following RH stroke was much higher than that reported by Biervoye et al.^40^ for their 11 subjects with stroke, were absent of clinical information, including lateralization of stroke, precluding any predictions of ToM specifically after RH stroke.

The ToM task was carefully selected to minimize confounds that have plagued prior research. However, the visual presentation of the task potentially could create a new confound for participants with unilateral neglect. Three participants were identified to have neglect. If the neglect impaired participants’ ability to fully process the visual task, errors would be expected across all conditions. While two of the participants identified with neglect also had deficits on the ToM tasks, the other did not evidence such deficits, suggesting that neglect in and of itself doesn’t necessarily prevent participants from processing the visual ToM task.

Clinically it will be important to determine the pattern and time course of recovery of ToM. If, like unilateral neglect, it recovers spontaneously in many cases, then clinicians can monitor change over time and treat as necessary if recovery is not detected. If, however, these deficits persist, early identification and treatment will reduce the overall impact on communication and socialization and long-term aspects of quality of life.

## Supplemental Information

**Supplemental Table 1.**
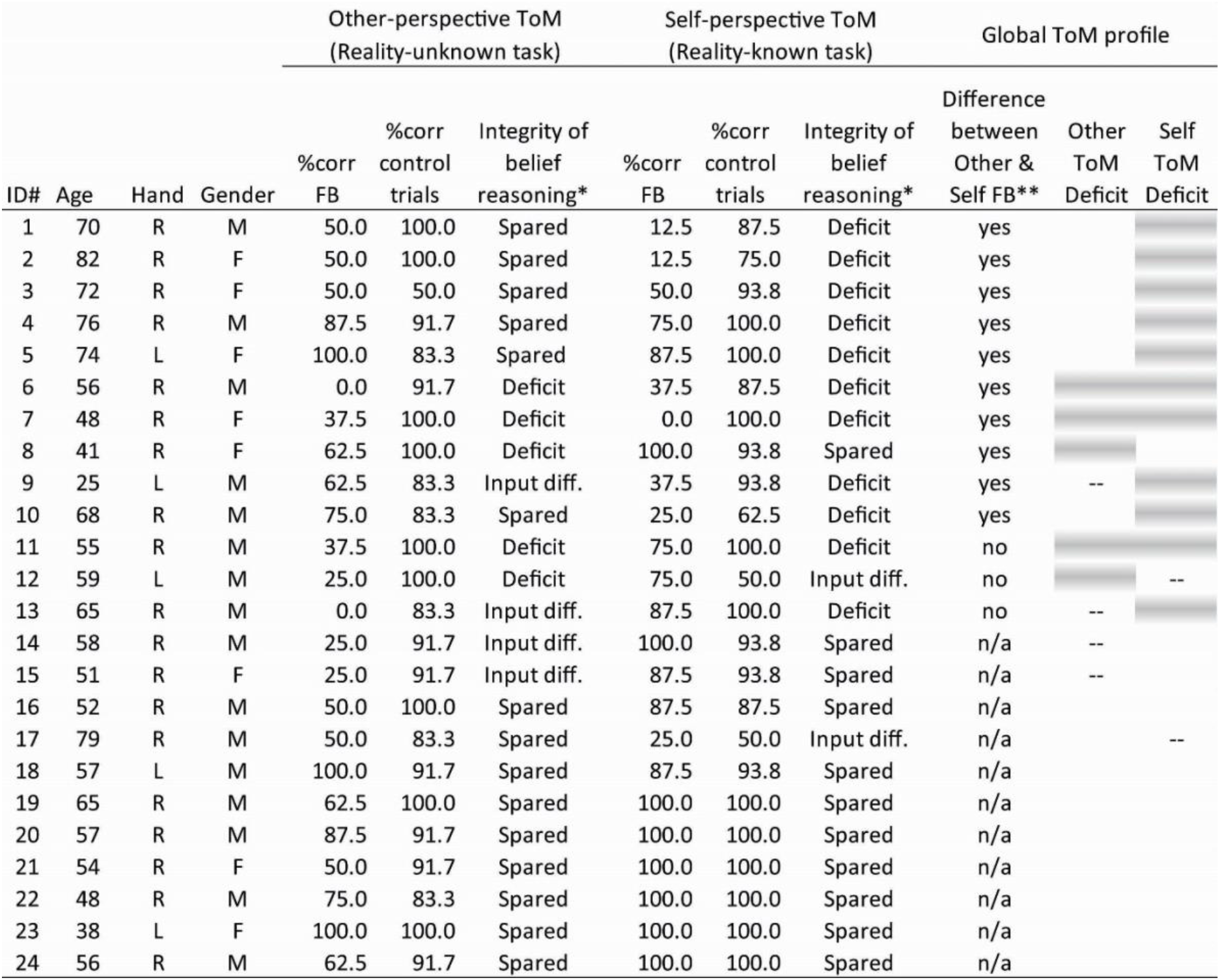
Participant demographics, % accuracy across Theory of Mind (ToM) conditions, and classification of ToM deficits based on age-matched control performance. % corr = % correct; FB = False Belief; input diff. = input processing difficulties; Shading = deficit; *Patient FB Z-scores < −2 SDs from non-brain damaged FB performance and control trial performance; **Difference between *Other* vs. *Self* ToM FB Z-scored performance > 2 SDs; -- False Belief performance affected by input processing difficulties.

## Footnotes

1 In the two cases when there was no variability in control performance (i.e. SD = 0), to calculate a patient’s z-score, we assumed SD = .2.

